# PSINDB: A comprehensive database of postsynaptic protein-protein interactions

**DOI:** 10.1101/2021.07.19.453019

**Authors:** Zsofia E. Kalman, Dániel Dudola, Bálint Mészáros, Zoltán Gáspári, Laszlo Dobson

## Abstract

The postsynaptic region is the receiving part of the synapse comprising thousands of proteins forming an elaborate and dynamically changing network indispensable for the molecular mechanisms behind fundamental phenomena such as learning and memory. Despite the growing amount of information about individual protein-protein interactions in this network, these data are mostly scattered in the literature or are stored in generic databases that are not designed to display aspects which are fundamental to understanding postsynaptic functions. To overcome these limitations we collected postsynaptic protein-protein interactions (PPIs) together with a high amount of detailed structural and biological information and launched a freely available resource, the Postsynaptic Interaction Database (PSINDB) to make these data and annotations accessible. PSINDB includes tens of thousands of binding regions together with structural features mediating and regulating the formation of PPIs, annotated with detailed experimental information about each interaction. PSINDB is expected to be useful for numerous aspects of molecular neurobiology research, from experiment design to network and systems biology-based modeling and analysis of changes in the protein network upon various stimuli. PSINDB is available at http://psindb.itk.ppke.hu/.

## Introduction

Synapses are communication points between neurons responsible for transducing information. Synapses can be broadly categorized into presynaptic sites releasing neurotransmitters, and postsynaptic sites taking in chemical signals. Cellular and organism level events, such as neuronal development, memory and long-term potentiation, strengthen synaptic connections [1]. Excitatory synapses contain a special morphological unit called postsynaptic density (PSD), that localizes directly under the membrane of the receiving cell [2]. The PSD is composed of thousands of highly conserved proteins; however the exact distribution and organization of components change during neuronal development leading to morphological differences in the PSDs of different brain regions [3]. Mutations in proteins of the PSD are responsible for severe neurological and psychiatric diseases [4], many of them likely to be heritable as suggested by a growing number of evidence, such as in the case of Autism Spectrum Disorder [5,6].

Protein-protein interactions (PPIs) play an essential role both in the maintenance of synaptic plasticity and in disease emergence. PSD proteins exhibit a high degree of multivalency presenting multiple binding sites stacked into single proteins. This multivalency aids the formation of an elaborate and dynamic network via highly modular architecture, provided by distinct structural and functional elements [7]. Intrinsically disordered regions (IDRs) were shown to be critical components for network assembly in the PSD [8], often by containing short linear motifs mediating transient interactions [9], post-translational modification sites regulating interactions as switches [10] and more. These regions play crucial roles in forming higher-order assemblies of the PSD, such as fuzzy complexes [11] and membraneless organelles via liquid-liquid phase separation [12].

Although there is a huge amount of information available about PPIs formed in the postsynapse, they are either scattered in the literature or they are collected in databases that were not prepared to store extended information that helps us to better understand the nature of synapse formation and function at the molecular level. Here we present the Postsynaptic Interaction Database (PSINDB, https://psindb.itk.ppke.hu/) aiming to provide a fuller and more complex picture of the postsynaptic protein network. PSINDB is a comprehensive database focused on postsynaptic proteins, containing a high number of collected, manually curated or derived interactions with precisely defined binding regions, enriched by various structural features of the constituting proteins. We believe that PSINDB will help scientists to get closer to the understanding of the complex nature of synaptic function.

## Methods

### Selection of postsynaptic proteins

We downloaded the human, mouse and rat reference proteomes (2021_April release) from UniProt [13]. We assigned orthologous proteins using the OMA database [14] and gene names. To define postsynaptic proteins the following sources were utilized: SynaptomeDB [15], Genes2Cognition [16], SynGo [17] and GeneOntology postsynapse/postsynaptic density terms (and their child terms) [18].

### Manual curation

For the first step in the curation we extensively searched PubMed and Google scholar for relevant publications. Since many of these publications were already included in other PPI databases, we only used articles that were not processed elsewhere. The curation work was done adhering to the HUPO-PSI Molecular Interaction (MI) workgroup standards, and at least MIMIx curation level [19] was used, while terms describing interactions were taken from the PSI-MI ontology (https://www.ebi.ac.uk/ols/ontologies/mi). We considered every described interaction between rat, mouse and human proteins, then mapped them back to the human reference proteome.

### Data integration

We integrated binary PPI information from the IntAct [20], BioGrid [21] and STRING [22] databases. Furthermore, we derived interaction data from PDB [23]: first we reconstructed the full structure using the BIOMT lines, then Voronota [24] was used to calculate intermolecular surfaces and to identify interacting residues.

We have predicted or downloaded the following protein structure-related information: transmembrane segments (Human Transmembrane Proteome database [25], liquid-liquid phase separation (PhaSePro [26]), short linear motifs (Eukaryotic Linear Motifs database, [27]), phosphorylation (UniProt annotations), domains (PFAM [28]), coiled-coils (Deepcoil [29] and UniProt annotations) disordered regions and disordered binding regions (IUPred2A [30]) and disease causing germline mutations classified in Disease Ontology [31]. Protein alignments between isoforms were done using ClustalOmega [32].

## Results

The homepage of PSINDB database is available at the URL:https://psindb.itk.ppke.hu/. We have created an interactive graphical user interface for the visualization of interaction data.

Two types of searches can be performed on the site I) the protein search will result in a list of proteins II) the interaction search will result in a list of interacting partners. In both cases the gene name, entrez and HUGO gene ID, UniProt ID and various aliases can be used as a query.

Clicking results from the protein search opens the individual protein pages, which includes a graphical scheme of the corresponding protein along with ten sections summarizing structural, interaction and network data (Figure 1). The sections are the as follows: 1) Evidence for PSD localization (with link to the source database) 2) Function (short description mirrored from UniProt); 3) Protein features (structural and interaction information, featuring interacting regions collecting binding regions of all partners); 4) Binary interactions with known binding regions (interacting regions with each partner one by one); 5) Network (a matrix-like representation of partners, where the depth of the color correlates with the strength of experimental evidence for the interaction); 6) Isoforms (alignment of isoforms highlighting segments not included in splice variants making it easily comparable with the location of the binding regions) 7) Disease-causing germline mutations (mutations and diseases associated to the protein, together with the partners where binding sites overlap with the mutated residue) 8) Linear motifs (together with the partners where binding sites overlap with the mutated residue) 9) Fingerprint (percentages of GeneOntology molecular function, GeneOntology biological process and disease ontology terms shared within the network of binary interaction partners of the protein) 10) All partners (including those not having postsynaptic annotations, yet the interaction indicate the partner may have the same localization).

**Figure 1:**
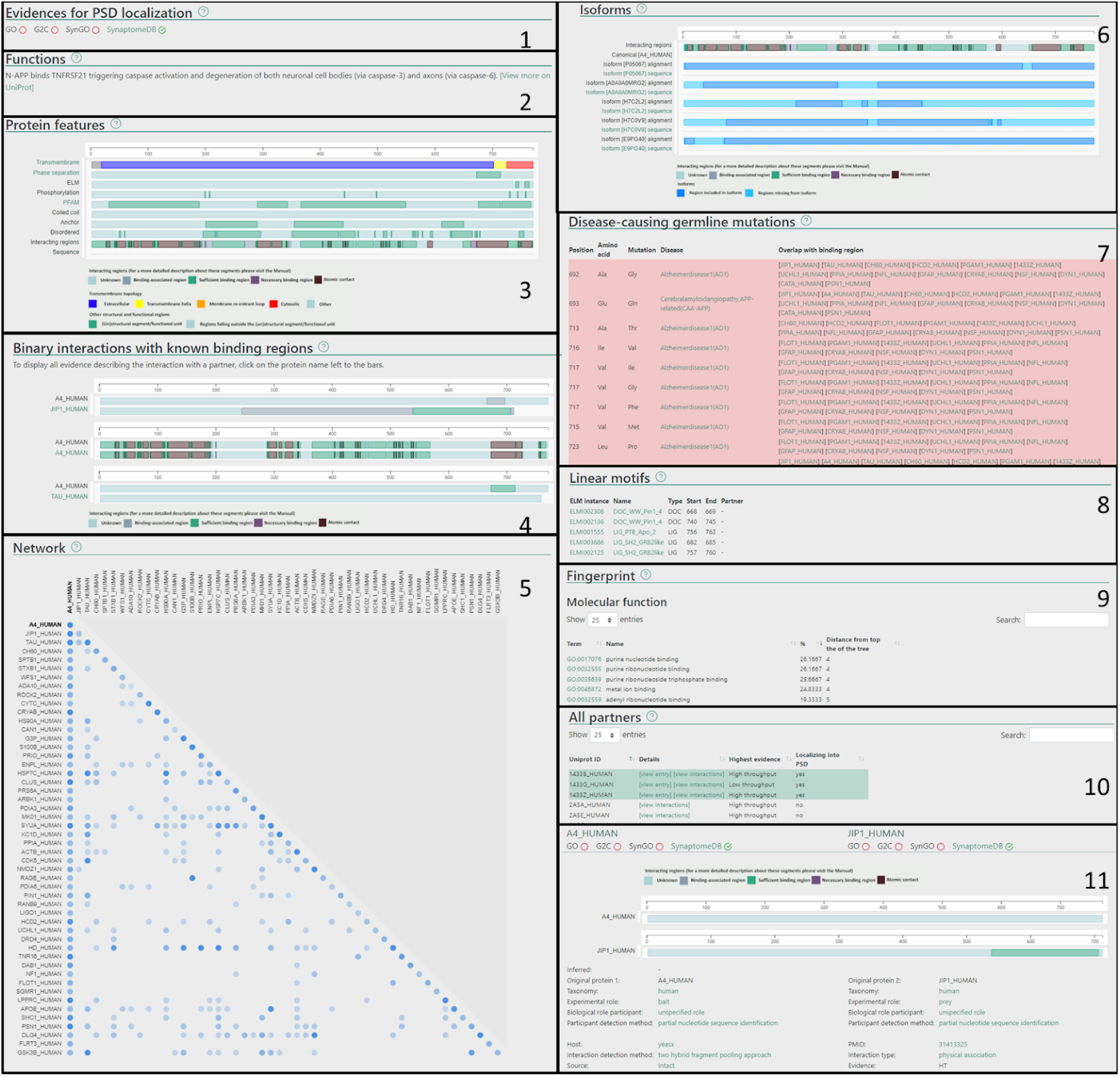
Layout of the PSINDB database with A4_human protein as an example. Protein page 1) Evidences for PSD localization 2) Function 3) Protein features 4) Binary partners with known interacting regions 5) Network 6) Isoforms 7) Disease causing germline mutations 8) Short linear motifs 9) Fingerprint 10) All partners. Interaction page 11) experimental details.

The interaction page can be accessed from the Protein page by clicking on the partner, or from the interaction search page. The interaction page contains the definition and details about the experiment (Figure 1), including, but not limited to the type of interaction, interaction detection method, host organism, experimental and biological role of participants and more. Binding regions for each experiment are also displayed where available.

In addition, we prepared several predefined subsets of postsynaptic proteins/interactions - including scaffolding proteins, proteins involved in phase separation, hub proteins and more - that may be interesting to a broad range of users.

PSINDB core data is currently available to download in the PSI-MI mitab format [33], with further download options planned in the near future.

## Conclusion

PSINDB is a comprehensive database of postsynaptic protein interactions, featuring detailed experimental information, binding regions of partners, and structural and biological annotations. The current version of PSINDB stores 16,618 unique binary interactions between 2,210 proteins, with 484,314 experimental evidence from 22,892 research articles. We expect that PSINDB will be a valuable resource for researchers investigating the molecular mechanisms of synaptic signal transduction, function, regulation and misregulation. In particular, our detailed data on protein-protein interaction sites can substantially help to understand how different complexes from the same components can be assembled and how these can interconvert into one another. PSINDB can serve as a resource for network and systems biology-based models of postsynaptic organization and function. PSINDB is available at https://psindb.itk.ppke.hu/

## Acknowledgement

Support from the National Research, Development and Innovation Office through grant OTKA 124363 to Z.G. is acknowledged. BM has received funding from the European Union’s Horizon 2020 research and innovation programme under the Marie Skłodowska-Curie grant agreement No. 842490 (MIMIC).

